# Clay larvae do not accurately measure biogeographic patterns in predation

**DOI:** 10.1101/2023.09.29.560167

**Authors:** Antonio Rodriguez-Campbell, Olivia Rahn, Mariana Chiuffo, Anna Hargreaves

## Abstract

**Aim:** Spatial variation in predation can shape geographic patterns in ecology and evolution, but testing how predation varies across ecosystems is challenging as differing species compositions and defensive adaptations can mask underlying patterns. Recently, biogeography has borrowed a tool from ecology –clay prey models. But clay models have not been adequately tested for geographic comparisons, and a well-known problem –that clay prey only appeal to a subset of potential predators– could lead to inaccurate detection of geographic patterns whenever the relative importance of predator guilds varies among sites. Here, we test whether clay larvae accurately capture geographic differences in predation on real larvae.

**Location:** 90° of latitude and >2000 m elevation across the Americas

**Taxon:** vertebrate and invertebrate predation on ‘superworms’ (*Zophobas* larve)

**Methods:** Across six sites that vary dramatically in latitude, elevation, and biome, we quantified predation on live, dead, and clay larvae. We physically excluded vertebrate predators from some larvae to distinguish total predation and invertebrate-only predation.

**Results:** Predation on live superworms almost doubled from our high-elevation high-latitude site to out low-elevation tropical site. Geographic patterns were highly consistent among live and dead larvae, but clay larvae missed extremely high predation at some sites and therefore mismeasured true geographic patterns. Clay larvae did a particularly bad job at capturing geographic patterns in predation by invertebrates.

**Main conclusions:** Clay larvae are inappropriate for large-scale tests of predation, and should be abandoned for biogeographic studies. Biogeographic experiments should instead employ realistic baits, and clay prey should be reserved for comparisons within, rather than across, predator communities.

## Introduction

The idea that species interactions vary geographically has intrigued ecologists for more than 150 years (Darwin, 1859). Stronger interactions have been proposed to explain the astounding diversity of tropical trees (pests and pathogens; Connell, 1971; Janzen, 1970), narrower elevational ranges in mainland vs. island systems (competition; MacArthur, 1972), and higher speciation rates in the tropics (all interactions; Dobzhansky, 1950; Schemske, 2009). Species-specific field studies have shown that local spatial variation in interaction strength drives biological patterns as diverse as plant mating-system evolution (pollination; Moeller, 2006), species longitudinal range limits (endophytes; Afkhami, McIntyre, & Strauss, 2014), and variation in sexual dimorphism (predation; Luyten & Liley, 1991). However, measuring the interaction intensity at larger geographic scales is more challenging, as it is difficult to control for regional variation in species composition and local evolutionary responses that can mask underlying geographic patterns (Rasmann & Agrawal, 2011).

An emerging approach to quantifying interaction intensity across large geographic scales is the standardized predation experiment, where experimenters set out the same standardized prey at a sites that span large changes in latitude, longitude, or elevation. Many standardized experiments have used real biological material, such as agricultural seeds (Hargreaves et al., 2019; Hillyer & Silman, 2010; Orrock et al., 2015), quail eggs (Loiselle & Hoppes, 1983; McKinnon et al., 2010), and insect larvae (Jeanne, 1979), which not only look but smell (and act, in some cases) like real prey items. While the realized ecological importance of an interaction will also depend on exposure time (Martin & Hargreaves, 2023) and trait variation (Chen, Hemmings, Chen, & Moles, 2017), large-scale standardized experiments have made major advances in quantifying geographic patterns in underlying predation rates, showing variation with latitude (Hargreaves et al., 2019; Jeanne, 1979; McKinnon et al., 2010), elevation (Camacho & Avilés, 2019; Hargreaves et al., 2019; Hillyer & Silman, 2010), and climate (Orrock et al., 2015; Rodriguez-Campbell, 2023). However, standardizing real prey items across an international experiment can be challenging, given difficulties in shipping biological material across international borders or finding the same material locally in different countries.

An alternate approach that is rapidly increasing in popularity is to construct artificial prey, often out of modelling clay (Lövei & Ferrante, 2017). Clay models facilitate standardization across countries (Roslin et al., 2017), and in theory enable researchers to distinguish among predator guilds based on attack marks left in clay (Howe, Lövei, & Nachman, 2009). However, up to half the attack marks (and all missing prey) cannot reliably be assigned to a predator group (Bateman, Fleming, & Wolfe, 2017; Rodriguez-Campbell, 2023; Rößler, Pröhl, & Lötters, 2018), and ~20% of mark identifcations are incorrect even when made by experienced scientists (Valdés□Correcher et al., 2022). Further, artificial prey lack important sensory signals like movement, sound, thermal and scent cues, that are used to varying degrees among and within predator guilds including snakes, lizards, spiders, and ants (Bernays, 1997; Cooper, 2008; Herzog Jr & Burghardt, 1974; Reilly & McBrayer, 2007; Young, 2003). Thus predation on clay prey is biased toward predators that use visual cues of colour, shape and contrast, such as passerine birds, jumping spiders, and mantids (Thery & Gomez, 2010). While this is recognized as a general limitation of clay models (Bateman et al., 2017; Rößler et al., 2018), its particular importance in a biogeographic context is not yet sufficiently appreciated.

If clay prey are biased in the predator guilds they measure, they will misrepresent geographic differences in predation rates when sites differ in the relative importance of predator guilds or foraging strategies. Empirical studies on real prey show conclusively that such geographic variation is common. For example, the relative importance of predation by ants decreases toward high elevations (Camacho & Avilés, 2019), snakes account for anywhere from 0 to 90% of nest predation at sites across North America (DeGregorio et al., 2014), and the relative abundance of avian foraging strategies varies among forests (Korňan, Holmes, Recher, Adamík, & Kropil, 2013). Biased detection should be especially problematic for prey that normally move, such as caterpillars and other larvae. Remarkably few studies have attempted to validate clay larvae against real larvae for comparing predation rates among sites (Greenop et al., 2019; Nagy, Schellhorn, & Zalucki, 2020), even though clay larvae are increasingly used for geographic comparisons (Gray et al., 2022; Roslin et al., 2017; Wallis et al., 2021; Zvereva et al., 2019).

Here, we test whether clay insect larvae accurately assess geographic differences in predation. We quantified predation rates on live larvae, dead larvae, and fake (clay) models of larvae at six sites that differed dramatically in latitude (-41° to 51°), elevation (69 to 2400 masl), and biome (forest, steppe, and alpine; Fig. 1A-G). Live prey should most closely reflect predation on wild prey; dead prey have realistic visual and scent cues but lack movement and behavioural cues; and clay prey have only coarsely realistic visual cues. We tested whether clay larvae detect the same biogeographic patterns in predation as real (i.e. live or dead) larvae. We did this for total predation (*Question 1*), and invertebrate-only predation (*Question 2*), since vertebrate and invertebrate predators differ in the sensory cues they use to find prey and their geographic patterns of predation (Rodriguez-Campbell, 2023). Rather than relying on attack mark identification, we physically excluded vertebrates from some larvae using wire cages (Fig. 1J).

**Figure 1.**
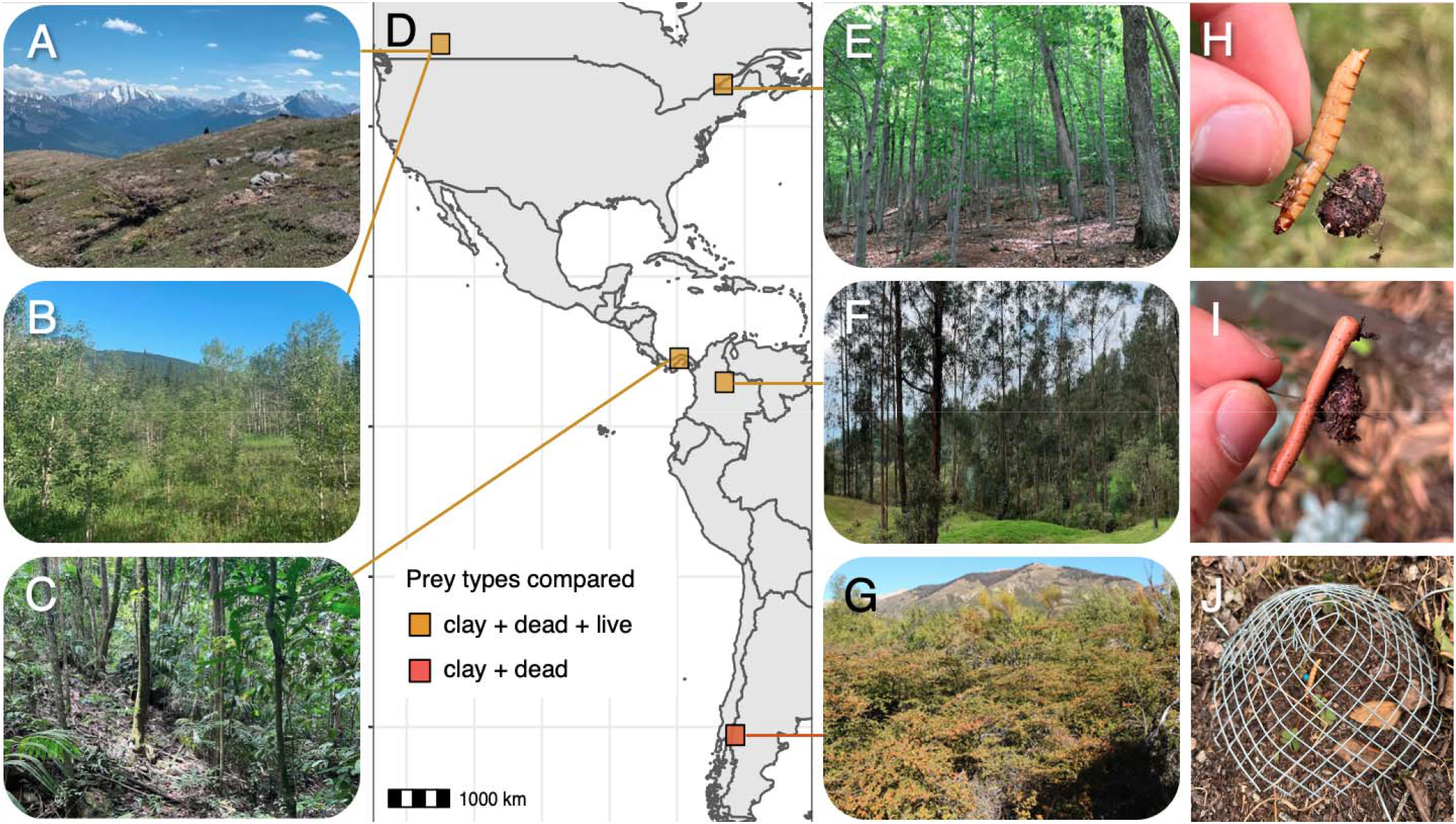
Experiment locations and setup. We measured predation at six sites: a high-elevation (A) and low elevation (B) site in Alberta, Canada; a temperate old growth forest in Quebec, Canada (E); a tropical rain forest in Panama (C); a tropical deciduous forest in Colombia (F); and a wet steppe shrubland in Patagonia, Argentina (G; site details in Table 1). We measured predation on real (live or dead) larvae of *Zophobas morio*, sold commercially as ‘superworms’ (H), and on clay replicas of superworms (I). To distinguish predation by invertebrates only, we excluded vertebrate predators from some worms using wire cages (J).

## Methods

### Study sites

We measured predation in 2021 and 2022 at 6 sites in the Americas (Fig. 1; Table 1). Three sites were in temperate North America: an aspen-boreal forest (1405 masl) and alpine meadow (2400 masl) in the Rocky Mountains of southern Alberta, and an old growth deciduous forest in southern Quebec (Gault Nature Reserve). Two sites were tropical: a rainforest in Panama (Gamboa), and a deciduous forest in central Colombia (near Duitama). One site was in temperate South America: a wet steppe shrubland in Patagonia, Argentina (near San Carlos de Bariloche; Table 1; Fig 2). At each site we set up our experiment at least 50 m from roads or trails.

**Table 1:** Site details and sample sizes. Each plot had 1 live, 1 dead, and 1 clay larvae. During each run we excluded vertebrate predators from some plots using wire cages to quantify invertebrate-only predation.

| Country | Region | Latitude | Elevation<br>(masl) | Biome | Runs | Total plots (replicates) |  |
| --- | --- | --- | --- | --- | --- | --- | --- |
|  |  |  |  |  |  | Total<br>predation | Invertebrate<br>predation |
| Canada | Alberta | 51.0 | 2400 | Alpine | 3 | 48 | 12 |
| Canada | Alberta | 51.0 | 1400 | Forest | 3 | 48 | 12 |
| Canada | Quebec | 45.6 | 280 | Forest | 3 | 48 | 11 |
| Panama | Colon | 9.1 | 70 | Forest | 2 | 36 | 4 |
| Colombia | Boyaca | 5.9 | 2990 | Forest | 3 | 48 | 7 |
| Argentina | Rio Negro | -41.1 | 860 | Steppe | 3 | 65 | 14 |

**Figure 2.**
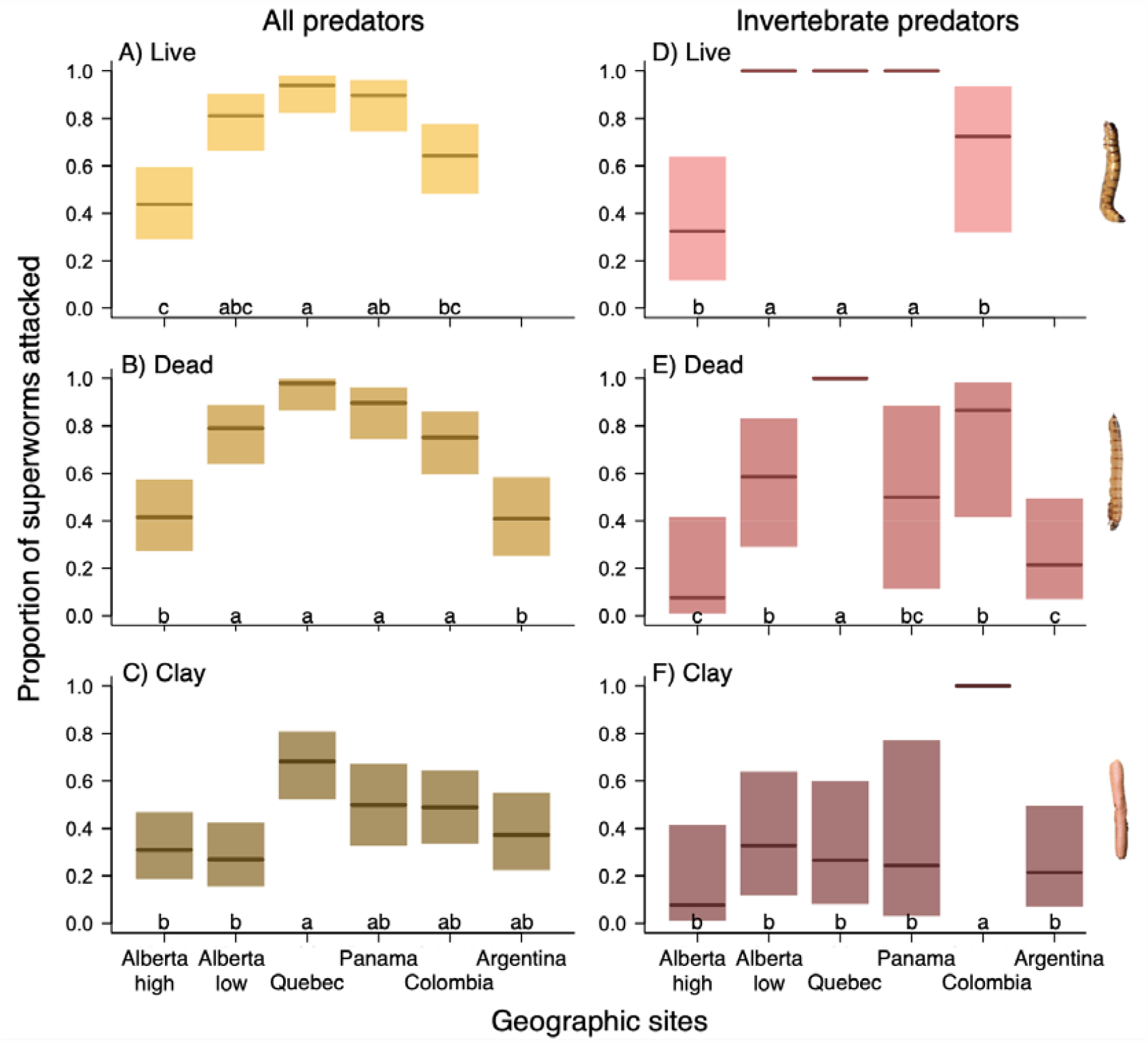
Clay larvae do not accurately detect geographic patterns in predation. This was true for both predation by vertebrate + invertebrate predators (A-C) and predation by invertebrate predators only (D-F; determined by caging larvae to exclude vertebrates). Thick lines and shaded bars show least squared means and 95% confidence intervals, respectively. Within each panel, sites that do not share any letters differ significantly in predation.

### Prey types

To standardize prey across countries, we chose a prey type widely available in petfood stores, known as superworms (Fig. 1). Superworms are the larvae of the large neotropical beetle species *Zophobas morio*, a member of the darkling beetle family (Tenebrionidae) native to Central and South America (Rumbos & Athanassiou, 2021). Superworms are large (5 cm long x 0.25 cm diameter) and active. They are sold as food for a wide variety of animals, including birds, reptiles, and mammals, making them an excellent candidate for comparing predation rates that might differ among predator guilds.

We compared predation rates on three types of larvae (Fig. 1H-J). To assess predation on prey with the full suite of natural sensory cues (colour, shape, smell, movement) we used live *Z. morio* larvae. To assess predation on prey with partial sensory cues (colour, shape, some smell but no movement), we used dead *Z. morio* larvae. Larvae were killed by freezing at -20°C 1 to 2 days before their use in an experiment (Nagy et al., 2020), and stored frozen until ~1 h before use. To assess predation on artificial prey models, which lack smell and movement cues but mimic the shape and colour of real larvae, we constructed artificial larvae from light brown, unscented FIMO modelling clay (5 cm long x 0.25 cm diameter).

### Experimental setup

On a given date at a given site, we selected ~20 to 30 locations (plots; Table 1), separated by ≥ 10 m from one another. In each plot we placed 1 live larva (all sites except Argentina), 1 dead larva and 1 clay larva, separated from each other by 1 m (Fig 1). We did not use live larvae in Argentina due to permitting issues. Above each larvae we tied flagging tape labelled with the plot number and larva type. We initially tried to tether superworms in place, but live worms buried themselves in soil. We therefore held them in place using a 5 cm sewing pin (i.e. too large to be ingested by small mammals and birds) (Hong, Held, & Williamson, 2011; Mathews, Bottrell, & Brown, 2011). To prevent live superworms from burying, we pinned each larva to a 1 cm ball of FIMO clay, then placed the clay ball in the top 1 cm of soil, held in place by the end of the pin. At our alpine site in Alberta, several pins were missing after the first run. As we did not wish to leave pins in the environment, we changed our method of securing worms in this site, instead using 1 cm bulldog clips to clip each larva to the end of a popsicle stick, which we stuck in the ground so that larvae were flush with the ground. Larvae stayed alive for the duration of the experiment, but died within 2 days of being pinned/clipped, thus there was no risk of superworms escaping into the environment. To better capture typical predation rates, we ran the experiment twice (Panama) or three times (other sites) per site, separated by at least 12 days (total = 17 runs).

To quantify predation by invertebrates without relying on attack mark identification, we caged larvae in 2 to 6 plots per run to exclude vertebrate predators (*Question 2*). Cages were cones of 1/2” mesh ~12 cm high x 15 cm diameter, secured tightly to the ground using metal lawn staples (Fig. 1J). We spaced caged plots evenly across each site.

We returned to record predation after 24 h. Each larva was scored as intact (no predation), or attacked (part of larva missing, bite marks or holes visible, or larva removed but pin and clay ball found). For clay larvae we attempted to assign attack marks to predator guild using an identification guide (Low, Sam, McArthur, Posa, & Hochuli, 2014), but ultimately we were not confident in the reliability of these scores so simply scored larvae as attacked or not. We excluded data from two plots (6 larvae) that we could not relocate. Our final sample sizes were 353 artificial larvae, 352 dead larvae, and 274 live larvae (fewer as live larvae were not used in Argentina), distributed among 293 uncaged plots (*Question 1*) and 60 caged plots (*Question 2*).

### Statistical analyses

We tested whether geographic patterns in predation intensity varied among prey types (live, dead, clay) using generalized linear mixed models (Bates, Mächler, Bolker, & Walker, 2014) in R v 4.2.0 (R Core team, 2020). R code and data will be publicly archived on acceptance. We used separate models for uncaged depots (total predation; *Question 1*) and caged depots (invertebrate predation; *Question 2*); a full model with a 3-way site x prey.type x predator.type interaction was unresolvable. Our response was binomial (was larva attacked or not, one data point per larva), so GLMMs used a binomial error distribution and logit-link function. Models included categorical fixed-effect predictors for prey type, site, and their interaction. As we did not set out live larvae at all sites, we analysed the data in two ways for each question. *Analysis 1* compared predation on live, dead and clay larvae at the five sites with live larvae. *Analysis 2* compared predation on dead vs clay larvae at all six sites.

The prey type x site interaction tests our core question of whether detected geographic patterns in predation (site differences) differ among prey types. We evaluated its significance using likelihood ratio tests comparing the model with vs. without the interaction compared to a χ ^2^ distribution. When the interaction was significant, we tested which sites differed significantly from each other for each prey type and vice versa using least squared mean contrasts, controlling for multiple comparisons (emmeans package; Lenth, Singmann, Love, Buerkner, & Herve, 2019). For invertebrate predation (*Question 2*), models struggled to converge or estimate realistic confidence intervals for contrasts, due to smaller sample sizes and multiple instances of 100% attack rates (Fig. 2). We therefore bootstrapped least squared mean contrasts 500 times (bootMer in lme4). Contrasts and least squared mean estimates were consistent between *Analysis 1* (all prey types) and *2* (all sites). We report both, but for visualization we plot back-transformed means and contrasts from *Analysis 1* for all sites but Argentina, and add results for Argentina derived from *Analysis 2*.

## Results

Attack rates on real larvae showed significant geographic variation among sites (Fig. 2). Predation by all predators (uncaged larvae) and by invertebrate predators (caged larvae) was extremely high at our lowland tropical site (Panama), but was also very high at our lowland temperate sites in Quebec and Alberta. Predation on real larvae was lowest at our alpine site in Alberta.

*Q1: Do clay and real prey detect the same geographic patterns in total predation?*

No – biogeographic patterns in predation by the full suite of possible predators (vertebrates + invertebrates) varied significantly among prey types (Fig. 2A-C). At the 5 sites with live, dead and clay larvae, live and dead larvae detected almost identical geographic patterns (Fig. 2AvsB), but clay larvae did not (Fig. 2C; prey type x site interaction: ^2^ = 20.4, *P* = 0.009). While clay larvae correctly identified the two sites with the highest total predation (Quebec and Panama), they experienced the lowest predation at the site that had the third highest predation on real prey (Alberta-low). These results are consistent when we compare just dead vs clay prey at all six sites (prey type x site interaction: χ ^2^ = 24.8, *P* < 0.001).

Uncaged clay prey were attacked significantly less often than real prey at four of our six sites, and experienced numerically (but not significantly) lower predation at the other two sites as well (Fig. S1). Predation on live and dead prey did not differ statistically at any site (Fig. S1).

*Q2: Do clay and real prey detect the same geographic patterns in predation by invertebrates?* No – biogeographic patterns in predation by invertebrates (caged larvae; Fig. 2D-F) varied significantly among prey types. At the five sites with live, dead and clay larvae (prey type x site interaction: χ ^2^ = 25.9, *P* = 0.001), live and dead larvae again detected similar geographic patterns, although patterns were not quite as consistent as for total predation (perhaps due to the smaller sample sizes for caged larvae). Clay models correctly identified the highest invertebrate predation site (Colombia), but missed high invertebrate predation in Panama, Quebec, and Alberta-low. These results are consistent when we compare just dead vs clay prey at all six sites (prey type x site interaction: χ ^2^ = 13.4, *P* = 0.020; Fig. 2E-F).

Predation by invertebrates differed more erratically among prey types than did total predation, although this may partly reflect our smaller sample size for invertebrate-only predation (Fig. S1). Live superworms experienced equivalent (3 sites) or higher (2 sites) invertebrate attack rates as dead worms. Invertebrate attack rates on clay worms were more erratic, being lower than on real worms at three sites, equivalent at two sites, and higher at one site (Colombia).

## Discussion

Predation rates on real larvae varied significantly among our six sites, but clay larvae did not capture the same geographic patterns in predation, neither for total predation nor predation by invertebrates only (Fig. 2). While clay larvae correctly identified the highest predation site for total and invertebrate predation, they missed extremely high predation levels at several sites. For example, clay prey missed the significantly higher predation at our low-elevation tropical site (Panama) compared to out high-elevation high-latitude site (Alberta-high), for both total and invertebrate predation. In contrast, live and dead larvae detected almost identical patterns in total predation (Fig. 2AvsB), and very similar patterns in invertebrate predation (Fig. 2DvsE). Thus the difference in clay models is not because predation rates are inherently hard to quantify consistently (which would be discouraging for large standardized studies!) but because clay models are not adequately detecting true underlying geographic patterns in predation. Clay larvae are thus they are an inappropriate tool for comparing predation intensity among sites.

Clay larvae seemed like an exciting method for biogeographic studies. They are cheap and easy to ship across international borders or to standardize even when made by people in different countries. They provide limitless opportunities to test the effects of prey size (Remmel & Tammaru, 2011), within-site location (Low, McArthur, & Hochuli, 2016), colouration and warning signals (Linnen et al., 2013; Noonan & Comeault, 2009; Remmel & Tammaru, 2011). Using clay models avoids any risk of site contamination and they are certainly easier to deploy than struggling live larvae (superworms are surprisingly strong). However, initial biogeographic studies glossed over the fact that the method had not been verified for geographic comparisons. We know that clay models lack important sensory cues, we know that predators differ in the sensory cues they use to hunt, and we know that the relative importance of predators varies among sites. Logic alone thus suggests that clay prey may therefore mis-detect site differences in predation rates, and here we confirm that suspicion with data.

How then should we interpret existing biogeographic studies based on clay larvae? Some of these studies have found convincing evidence for geographic patterns that have also been found using real biological prey. For example, Roslin et al (2017) found that attack rates on clay caterpillars declined strongly toward high latitudes and elevations; findings consistent with longstanding theory (Darwin, 1859) that are supported by large-scale standardized experiments assessing predation on live wasp larvae (Jeanne, 1979) and real seeds (Hargreaves et al., 2019). We suggest that publication bias may play a role; studies based on clay larvae are more likely to be published in high-profile journals when their results demonstrate an expected biogeographic pattern, and the clay larvae method more likely to be questioned when results are less clear or predicted.

Could clay models still be useful if we conduct pilot studies to verify that they detect similar patterns as real prey? Our results suggest not. Clay prey can yield accurate geographic comparisons for some sites (e.g. Alberta-high vs Quebec) while still missing huge differences among other sites (e.g. Alberta-high vs Panama; Fig. 2A-C). Similarly, studies comparing predation on clay vs real prey within sites find that clay prey are good indicators in some contexts at some sites (Low et al., 2016) and poor indicators in others (Greenop et al., 2019; Nagy et al., 2020). Clay prey would therefore have to be validated against real prey for the full suite of intended geographic sites, at which point one might as well just use the results from real prey.

### Conclusions and Recommendations

Large-scale standardized experiments offer immense power to test important biogeographic patterns. Below, we make three recommendations to improve their use in biogeography.

1. Clay models should be abandoned for comparing predation rates among sites. A possible exception is when studies target an appropriate predator group (e.g. insectivorous birds), as long as patterns are corroborated with a method that does not rely on clay prey and attack mark identification (e.g. camera traps) and the limits to interpretation are made clear (e.g. González-Gómez, Estades, & Simonetti, 2006; Sam, Koane, & Novotny, 2015). It is not enough to admit that clay models yield biased results in the supplementary material but make broad claims in the Abstract.
2. Predation experiments using standardized prey should use the most realistic bait possible. Real items (eggs, seeds, live larvae) give the most confidence inspiring results. For mobile prey, dead prey may be a good compromise – they are scented and edible, detect similar predation rates to live prey (Fig. 2), but do not need to be secured in place. Another creative option is to use edible prey models, such as flour-lard caterpillars (Remmel & Tammaru, 2011; Sam, Remmel, & Molleman, 2015) or artificial seeds made of ground seed (Brehm, Mortelliti, Maynard, & Zydlewski, 2019).
3. Clay prey can still be useful for assessing predation differences *within* sites. However caution should be used even at small geographic scales whenever the relative importance of predator guilds is like to vary, including comparisons among habitat types or seasons (Nagy et al., 2020). For example, the increase in predation from our high-elevation Alberta site to our low-elevation Alberta site (separated by <1 km) rivaled the increase we saw from our high-elevation Alberta site to our high-elevation site in tropical Colombia, separated by more than 6000 km of latitude (Fig. 2).

One of the key challenges in biogeography is to experimentally test its hypotheses. We should continue to seek novel approaches to test biogeographic patterns across disparate sites, but we should also discard approaches once we realize they do not do so accurately.

## Acknowledgments

Sincere thanks to Sarah Chaddock and Anika Anderson for help wrangling wriggling superworms in the field. We are also grateful for funding from the National Sciences and engineering Research Council of Canada (NSERC) in a Discovery grant to ALH and scholarship to OR, to MITACs GlobalLink internship which helped fund ARC’s work in Colombia and Panama, and to Pablo Stevenson for helping host ARC while in Colombia.

## Appendix 1

**Figure S1.**
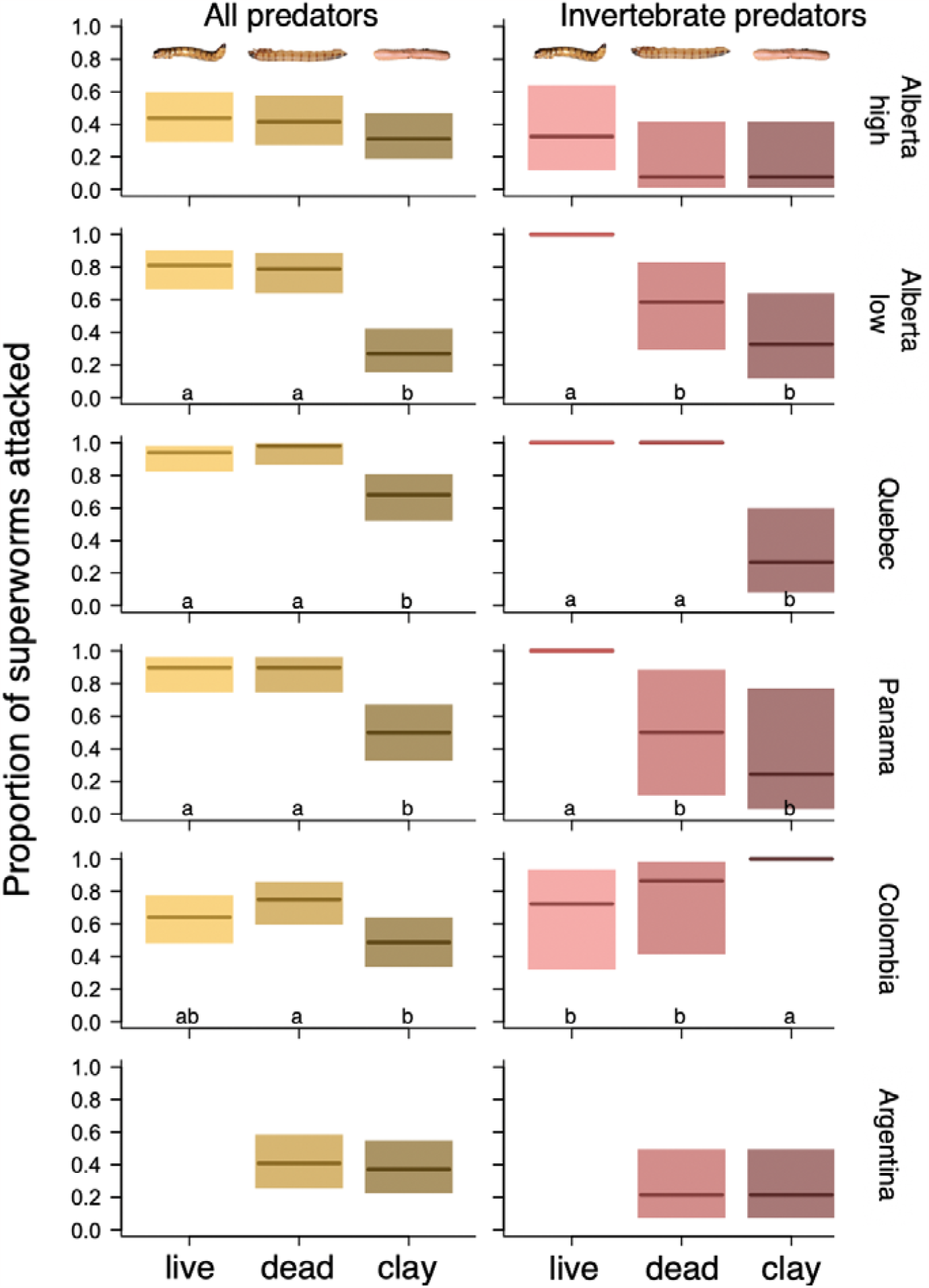
Predation rates on live, dead, and clay ‘superworm’ larvae. Total predation (left panels) was generally consistent between live and dead larvae, but often lower for clay larvae. Predation by invertebrates only (right panels; measured by physically excluding vertebrates from some larvae using wire cages) varied less consistently among prey types, perhaps due to smaller sample sizes for caged larvae. Thick lines and shaded bars show least squared means and 95% confidence intervals, respectively. Within each panel, prey that do not share any letters differ significantly in predation (in panels with no letters, no prey types differed significantly).

